# D-graph clusters flaviviruses and β-coronaviruses according to their hosts, disease type and human cell receptors

**DOI:** 10.1101/2020.08.13.249649

**Authors:** Benjamin A. Braun, Catherine H. Schein, Werner Braun

## Abstract

**Motivation:** There is a need for rapid and easy to use, alignment free methods to cluster large groups of protein sequence data. Commonly used phylogenetic trees based on alignments can be used to visualize only a limited number of protein sequences. DGraph, introduced here, is a dynamic programming application developed to generate 2D-maps based on similarity scores for sequences. The program automatically calculates and graphically displays property distance (PD) scores based on physico-chemical property (PCP) similarities from an unaligned list of FASTA files. Such “PD-graphs” show the interrelatedness of the sequences, whereby clusters can reveal deeper connectivities.

**Results:** PD-Graphs generated for flavivirus (FV), enterovirus (EV), and coronavirus (CoV) sequences from complete polyproteins or individual proteins are consistent with biological data on vector types, hosts, cellular receptors and disease phenotypes. PD-graphs separate the tick- from the mosquito-borne FV, clusters viruses that infect bats, camels, seabirds and humans separately and the clusters correlate with disease phenotype. The PD method segregates the β-CoV spike proteins of SARS, SARS-CoV-2, and MERS sequences from other human pathogenic CoV, with clustering consistent with cellular receptor usage. The graphs also suggest evolutionary relationships that may be difficult to determine with conventional bootstrapping methods that require postulating an ancestral sequence.

**Availability and implementation:** DGraph is written in Java, compatible with the Java 5 runtime or newer. Source code and executable is available from the GitHub website (https://github.com/bjmnbraun/DGraph/releases). Documentation for installation and use of the software is available from the Readme.md file at (https://github.com/bjmnbraun/DGraph).

**Supplementary information:** Supplementary information Table S1 and Fig. S1 are online available.

## 1 Introduction

While there are many methods for analyzing groups of distantly related sequences, most rely on sequence alignments to suggest phylogenies. Phylogenetic trees based on such alignments imply linearity and a common root, and become difficult to interpret as the number of sequences increases. The limitations of these methods become obvious when determining relationships among the thousands of viral sequences now available. Linear “distance trees” generated from pairwise alignments cannot reliably suggest evolutionary relationships between distantly related species, or correlate sequences according to disease phenotype or host. Authors typically resort to drawing 2D-plots by hand to illustrate the interrelatedness of larger virus groups.

Here, we present a rapid graphical method for analyzing large data sets of related protein sequences that does not require pre-alignment or assumption of a common ancestor. D-graph can present conventional pairwise alignment scores, such as those from Clustal, or simple overall identity. However, the program’s ability to generate “property distance” PD-graphs, based on physicochemical properties (PCP) of the amino acids (1), allows it to suggest more meaningful relationships among distantly related sequences. We have previously validated the PD method as a way to classify allergenic proteins and detect similar IgE epitopes (2, 3). We have shown that changes in the PCP values of key positions within flaviviral protein sequences correlate with significant phenotypic changes (4, 5). We have previously shown how PD-graphs clarify the inter-relationships of allergenic proteins (6) and group alphavirus and related PCP-consensus proteins (7).

In addition to describing the program, we show here its application to three diverse families of positive strand RNA viruses, flaviviruses (FV), enteroviruses (EV) and the β-coronaviruses (β-CoV), which include SARS, MERS and the pandemic SARS-CoV-2). The results illustrate how PD-graphs of the viral sequences correlate with phenotype and suggest evolutionary relationships of distantly related viruses.

## 2 Material and Methods

### 2.1 The property distance (PD)

The peptide similarity search tool (2, 3, 8, 9) was initially developed to find protein sequences in the Structural Database of Allergenic Proteins (SDAP) (10) containing user specified peptide sequences. The search tool uses a novel technique to find similar sequences in the proteins by comparing the physicochemical properties (PCPs) of the amino acids in the query and the target sequence. The differences in the PCP values in the two sequences are then measured by a property distance (PD). Briefly, five quantitative descriptors of physicochemical properties (PCPs) are assigned to each of the 20 amino acids. The five descriptors E1 to E5 were derived by multidimensional scaling of 237 physical–chemical properties for the 20 naturally occurring amino acids, thus the main differences of all 237 properties for the 20 amino acids are reflected by the five descriptors E1–E5. These in turn represent groupings of PCPs such as hydrophobicity, size, or secondary structure propensities, charge, aromaticity and size. The PD of two sequences A and B is then calculated as the average distance between the descriptor vectors E for corresponding amino acids, *i*.*e*.:

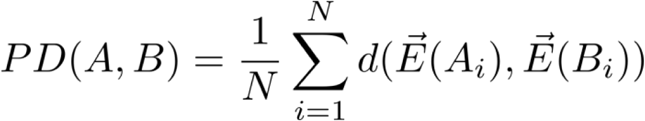

where *d* computes the standard Euclidean or L2 distance and *N* is the length of sequence B, assumed to be the same as A in this equation (Section 3.1 discusses the non-equal-length case.)

Identical sequences have a PD value of 0. Small PD values up to 4 typically indicate few substitutions with smaller values for conservative substitutions between the two sequences. Thus, the lower the PD value, the more closely related the sequences. In a database search of over 1500 allergenic protein sequences in SDAP, PD values of less than 8 are statistically significant for windows up to 12 amino acids and have been shown to correlate with immune recognition (2). Additional statistical measures (z-scores) can be calculated to indicate the significance of a PD value comparing it to the distribution of PD values over all random matches using larger datasets.

### 2.2 The DGraph program

DGraph, as discussed below, can generate a 2D map based on any input value list. In default mode, if given a list of FASTA formatted sequences in a text file as input, it calculates the pairwise PD values for the sequences and graphically presents their similarities. The resulting “PD-graph” represents the sequences as nodes in a 2D mapping where the distances of the edges between nodes are fitted to the numerical PD values. Whether the values come from the internally calculated PD values, or other user supplied ones, D-Graph calculates a metric of how faithfully the distances between all pairs of nodes on the graphic match their corresponding similarity scores or PD values.

### 2.3 Calculating PD values with a sliding window

For comparing sequences of amino acids that are not matched one-to-one, we instead define a metric comparing shorter fixed-length subsequences, or “windows.” We define the windowed PD between two sequences to be the PD between the least distant pair of subsequence having some length wSize (for the window size). The windowed PD measures the sequence similarity of the most conserved portion of the two sequences of length wSize. Note that windowed PD is 0 between two sequences having an identical string of amino acids at a conserved region of length wSize. The occurrence of this is exponentially less likely as wSize increases; for the results in this paper we use a wSize of 22 amino acids. Windowed PD can be computed naively by considering each pair of wSize subsequences in turn and computing their PD. A more efficient approach exploiting the linearity of PD is to slide a window of length wSize along and compute PD incrementally, for each offset of the shorter sequence along the longer one, and keeping track of the smallest PD value found.

### 2.4 Algorithm to find the optimal configuration of the nodes

Similarity scores from alignment programs, such as ClustalW, MUSCLE or T-Coffee are translated into distances as:

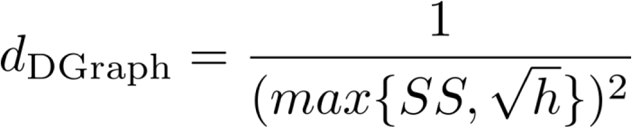

where *SS* is a similarity score and parameter *h* determines the minimum similarity cutoff. Similarity scores below *h*^0.5 are mapped to the maximum distance of 1/*h* in this equation, while larger similarity scores have an inverse square law with distance. The squaring reduces distances between highly similar sequences, which encourages visual clustering of similar sequences in the final figure DGraph produces.

The distances derived from the scores or the PD values are normalized and then used to compute a measure *U* of how accurately the represented distance between nodes matches the normalized distances:

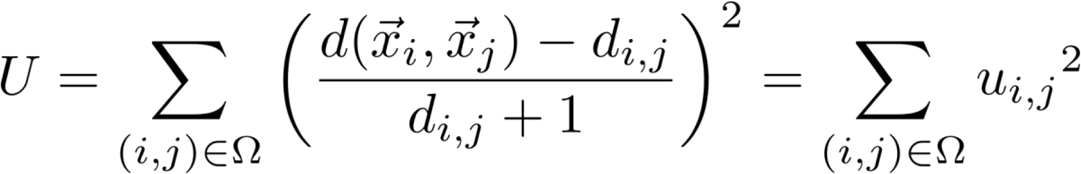

where the sum is over the set Omega of pairs of distinct sequences i,j, and the terms in the numerator are the representation distance and the normalized distance between those two, respectively. *U* is a sum of the squared relative error of each pair of sequence’s representation placement vs the normalized sequence distance, where relative error (*u*_*i,j*_) is computed with an adjusted denominator of *d*_*i,j*_ + 1 to account for 0-distance pairs.

As described, optimizing *U* results in a figure determined primarily by the most distant pairs of sequences. To better represent relationships between closely related sequences, we optimize a distance-adjusted *U\**, as follows: Define *u*\*_i,j_, equal to *u*_*i,j*_, but divided by d(*x*_*i*_, *x*_*j*_), represent a distance only when the representation distance is at least 0.001% of the figure diameter, and *U\** = sum(*u\**_*i,j*_^2^). Additionally, we remove from Omega any pair *(i,j)* with a PD or user-defined score-based distance larger than a configurable maximum distance comparison cutoff (the default value is 14). Sequences with no distance below this cutoff to the rest of the figure then become “islands” that are removed before optimization begins. These steps are both intended to make the resulting DGraph figure more faithfully represent short distances.

Initially, DGraph creates randomly-placed nodes for each sequence. DGraph then minimizes *U\** using a gradient descent approach. That is, at each step each node’s position *x*_*i*_ is shifted by an amount proportional to -*u\**_*i,j*_ along each direction (*x*_*j*_ – *x*_*i*_). In order to damp oscillations and promote convergence, we add a momentum vector *p*_*i*_ to each node, and apply the contribution of each *u\**_*i,j*_ to the momentum vectors rather than the position directly. Thus, at each step the position of each node *x*_*i*_ and its associated momentum vector *p*_*i*_ are updated as follows:

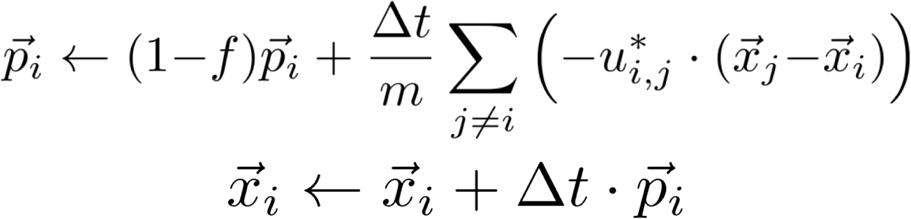

where small positive coefficients time-step (delta *t*), mass (*m*), and friction (*f*) are parameters of the descent.

### 2.5. Use of the program and utility tools

Parameters of the optimization, such as time-step, mass, friction, and maximum distance comparison cutoff, can be user specified with default values of 0.001, 0.01, 0.008, and 14. DGraph can be run interactively in order to fine-tune these parameters and to see a live view of the optimization. DGraph can be run with a helper script that runs the optimization multiple times, potentially resulting in different figures. The script compares the runs by optimization score, and normalizes the orientation of the final locations of the nodes (by rotation and reflection) to produce a consensus graphic (this uses the Kabsch algorithm, for details see (11)). The user can customize how sequences are labeled on the generated graphic, such as including the whole of short sequences (when graphing peptides, see (6)) or FASTA names as labels. The user can also apply custom coloring to the graphic including a color gradient for the line segments between nodes based on PD-value or user defined distance, and files of per-node colors useful for annotating sequence properties such as phenotype.

## 3. Results

### 3.1 DGraph converts various sequence scoring functions or PD values into 2D maps

In previous studies we established and validated the correlation of PD values for peptide segments with IgE binding affinities (2). We therefore expect that PD-graphs would be also useful to predict antibody cross-reactivities of different viral species. In supplementary Fig. S1 we illustrate this feature for the PCP-consensus sequence 7P8 of the domain 3 of the E protein of the four DENV viruses. The consensus domain 7P8 was designed as a potential vaccine candidate against all four DENV species and was recognized by all four species. In the PD-graph 7P8 is located near the middle of all four viruses (Fig. S1).

An alignment of 49 different FV was then used to illustrate the flexibility of the program. Figure 1 illustrates the flexibility of the software, showing 2D-maps for three different sets of similarity data for FV sequences based on different metrics. The first column shows a D-Graph calculated from previously calculated Clustal W alignment scores. The other two columns show two different ways to use PD values as a metric for generating the maps. The middle column shows the result of computing the PD value between each pair of sequences after a multiple alignment, which causes each sequence to be the same length by inserting alignment gaps. The last column shows results using instead a “sliding window” of 22 amino acids, by sliding every window in the shorter sequence along the longer sequence to find a best match, i.e. the one with the lowest pairwise PD-value, and computing the average PD-values of the matches.

**Fig. 1.**
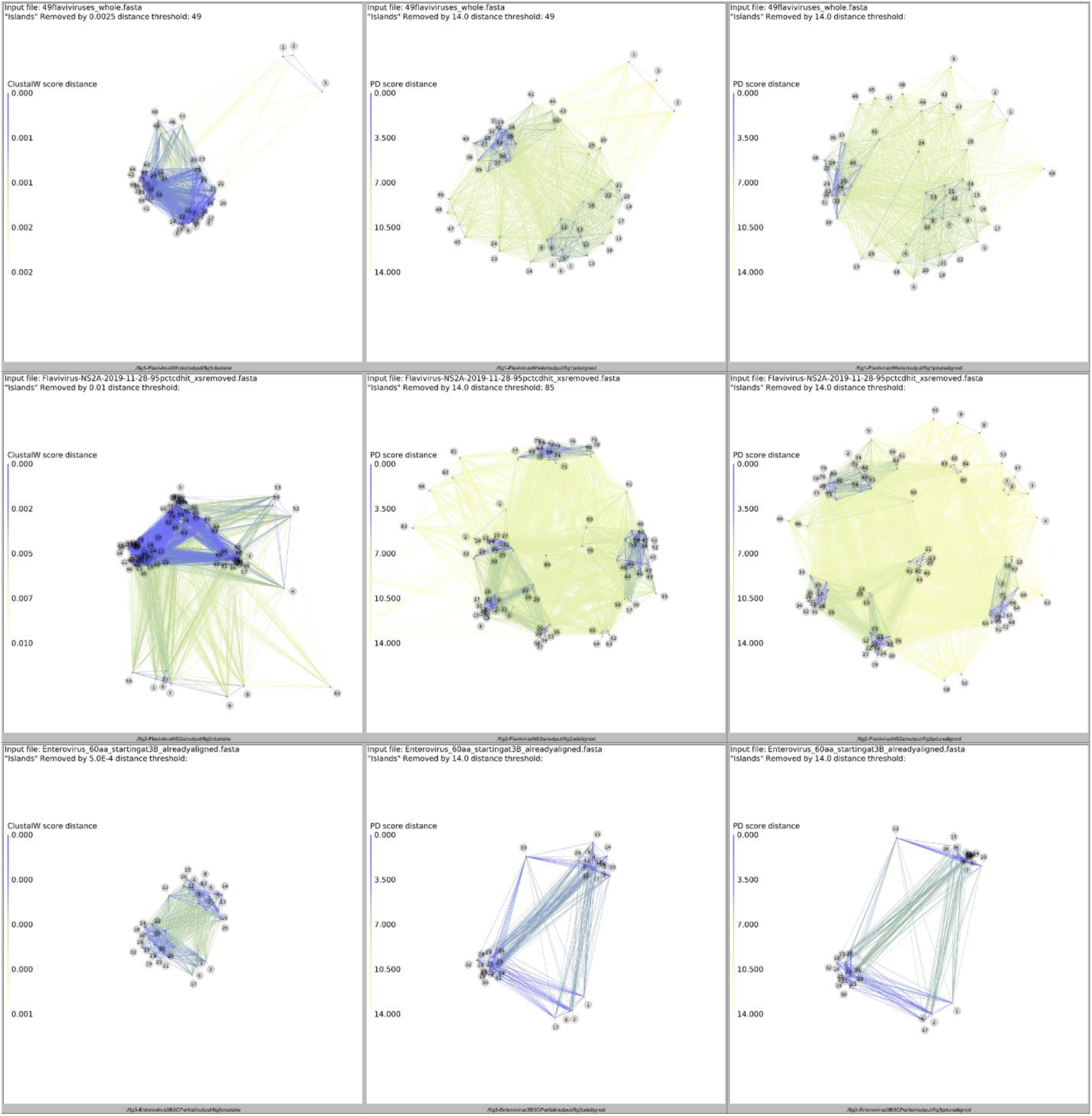
**Top row**: Screen shots of the output of DGraph for the analysis of the 49 polyproteins of 49 flaviviruses (Table S1) with the ClustalW score (left panel), and the PD values of 22_residue windows with aligned (middle panel) and unaligned sequences (right panel) **Middle row**: Screen shots from DGraph for the analysis of 86 sequences of the NS2a protein (all ∼234 aa long) of various FV, based on ClustalW scores and their pairwise PD values. The sequences were automatically downloaded from a Blast search, identical sequences removed and the resulting FASTA files subjected to Clustal W analysis or our PD based method with minimal involvement (except that the sequence headers were manually shortened and the sequences inspected to remove fragments). The resulting maps, which are similar to how PFAM B families are generated, show how PD-maps show a finer distinction among the viruses than simple Clustal scores using an unsupervised alignment. **Bottom row**: Screen shots from DGraph for the clustering of enteroviruses. A 60 amino acid sequence around the VPg protein (22 amino acids) of human and a few animal enteroviruses was used as input. The 2D plots clearly separate the simian viruses from the human ones.

The bottom frames of the figure show the results from a 60 amino acid region of the enterovirus 3B-3C protein interface, covering the viral protein linked to the genome (VPg) and an area in 3C that contains a vestigial additional VPg-like sequence (12, 13). For all three sets of viral sequences, the maps generated by the PD values show more distinct groupings than the maps generated by the ClustalW scores. They illustrate that the program can be used to rationally cluster even very long sequences (the whole viral genomes of the FV) as well as short sequences (the VPg area of the enteroviruses).

### 3.2 The PD-graphs correlates with vector and host competence

The map from the top row (middle panel) in Fig.1 is further annotated and highlighted in more detail in Fig. 2. The PD metrics shows a clear division of the tick from the mosquito borne or no-known vector and Rio Bravo group viruses. The map also clusters viruses that infect bats, camels (Kadam) and seabirds separately from those infecting humans, such as the Greek goat (node 29) and Turkish sheep encephalitis viruses (node 30) group. This observation also holds when the analysis is based on a smaller section of the virus, the 230 amino acid NS2a protein (middle row of Fig. 1). The PD-graph is consistent with the patterns of insertions and deletions we previously noted within the E-protein that mark the flaviviruses FV (4) according to species and disease specificity. With regard to the latter, hemorrhagic Yellow Fever (YF) virus (node 26) lies near the 4 DENV serotypes (nodes 19-22), which can cause hemorrhagic disease with fatal consequences in children. Interestingly, Zika virus (node 18) falls exactly between the mosquito borne encephalitic and hemorrhagic groups, consistent with its cross reactivity with DENV antibodies (14). An identity matrix for the E protein domain 3 region of Zika compared to WNV, DENV strains and our 7p8 PCP-consensus protein that binds antibodies to all four DENV serotypes (15) illustrates how this Zika indeed lies between the encephalitic and hemorrhagic FV, with >50% identity to all (Supplementary Fig. 1). While Zika infections generally cause mild disease, they can also result in Guillame-Barre syndrome and microcephaly if contacted by a pregnant mother. Zika’s nearest neighbor in the plot is the encephalitic virus, Rocio (16).

**Figure 2.**
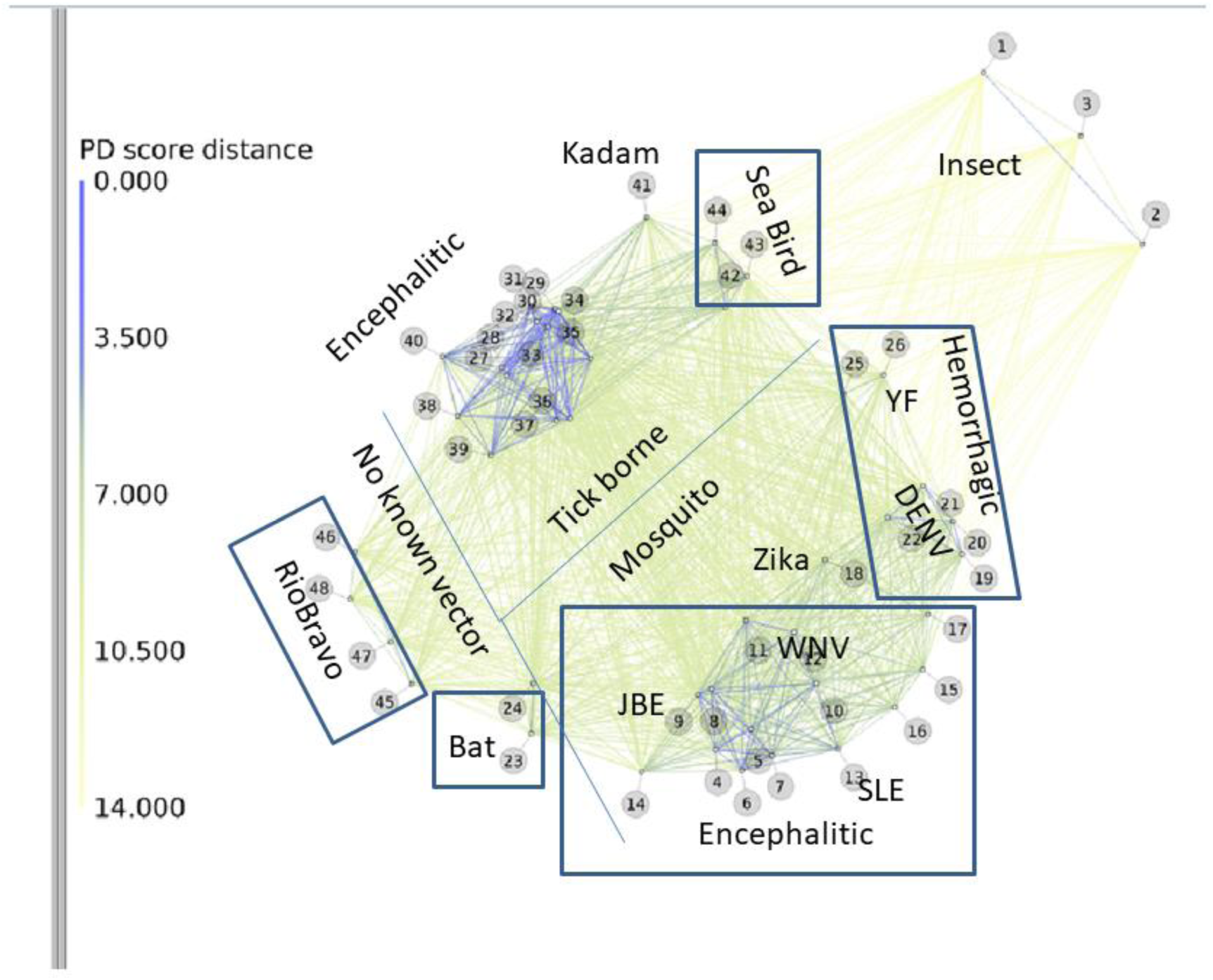
Annotated PD-graph of Flavivirus phylogeny from the polyproteins of 49 viral species. Each number in the plot corresponds to the viruses listed in Table 1 with FlaviTrack ID, Genbank, species name and length of amino acids. Starting from the unaligned FASTA sequences of the polyproteins, the program calculated pairwise PD values of all corresponding 22-residue segments for the polyproteins. The average PD values of the windows was used as a metric for the similarity of each pair. Lines in the figure indicate the degree of relatedness of the sequences, with a color code from blue to green as indicated with the PD values on the left of the figure. Blue thick lines indicate highly related flaviviruses. Divider lines and boxes were drawn by hand to emphasize the phenotypic groupings of the viral species.

The enteroviruses (EV), a non-enveloped group of +-strand RNA viruses that includes poliovirus, coxsackie and many other human pathogens, do not depend on arthropod vectors for transmission. The bottom part of Fig. 1 and the annotated graph (Fig. 3) show that even a short segment of EV sequences was sufficient to separate simian from human isolates. The separation by the two parameters (Fig. 1 g-i), Clustal or PD, generated similar clusters of the human viruses, which are consistent with other genetic classifications done with other areas of the sequences (17). Classifying enteroviruses according to disease type has proved very difficult and will not be attempted here. Early attempts to distinguish distinct types according to paralysis characteristics observed in a murine model, proved unreliable (18). Thus sequence similarity has proved to be the best classifier for strains,

**FIGURE 3.**
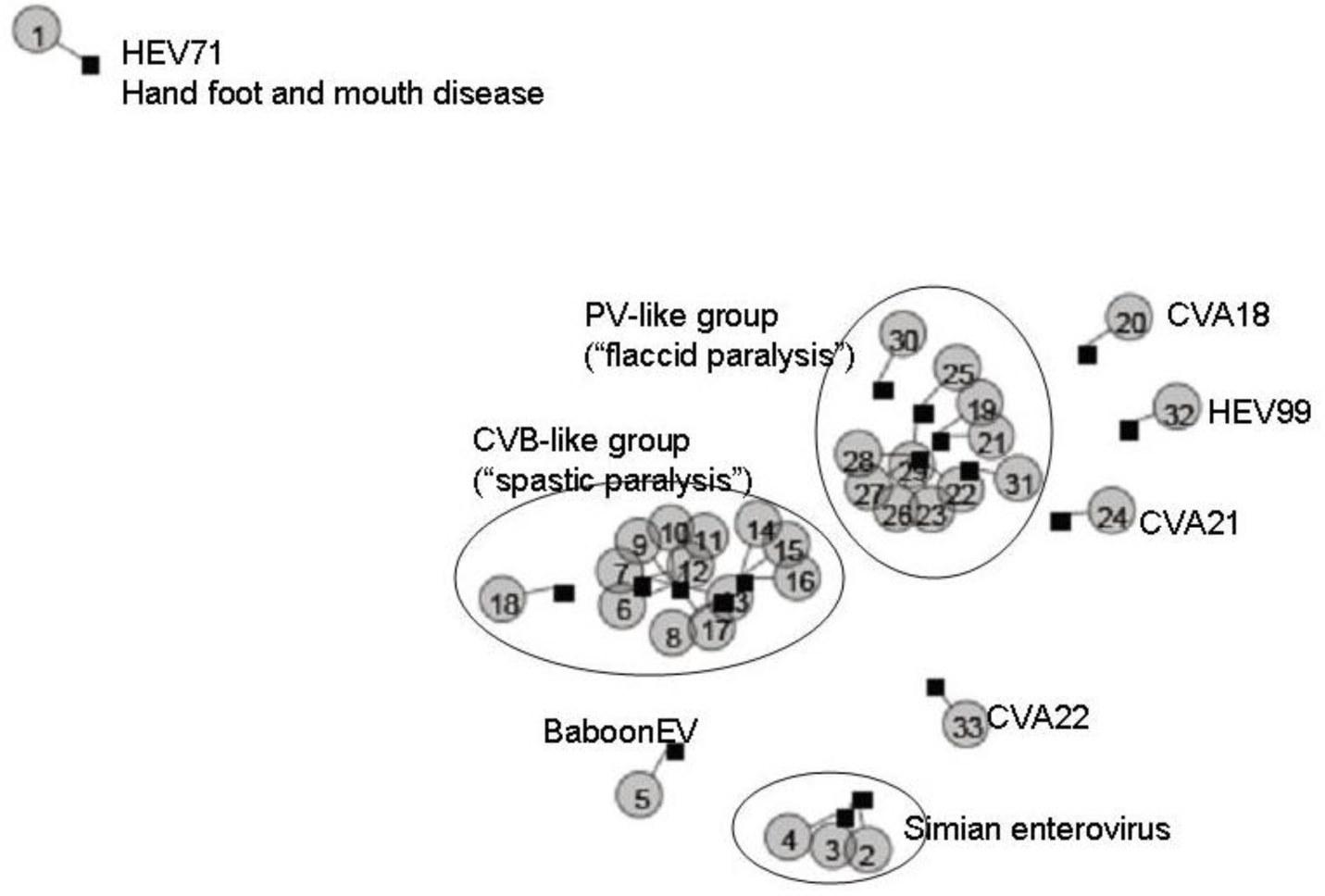
Annotated 2D-map for the enteroviruses based only on a short region encompassing their VPg sequences and part of the 3C protein. The clustering clearly separates the simian and baboon from the human viruses. The separation of the the PV-like group (“enterovirus type C”) from those resembling the coxsackie virus B group is based solely on sequence, as designation according to paralysis type has not proved reliable.

### 3.3 PD-graph accurately separate β-CoV according to disease type and receptor used

SARS-CoV-2 is known to be closely related, in its sequence, structure (19), receptor binding (20) and epitopes recognized by neutralizing antibodies isolated from survivors (21-23) to the SARS-CoV-1 virus that caused many deaths in a brief epidemic that ended in Asia in 2003 (24). It is more distantly related to the lethal Middle East respiratory syndrome coronavirus (MERS) (25). As an additional test of the program, 314 sequences of the Envelope 2 protein for diverse β-CoV were downloaded from the ViPR database (in March, 2020) and a single unaligned Fasta file used as input to the program. As Figure 4 shows, the resulting PD-graph, using the 22 amino acid window method, cleanly separates the SARS virus from the 2002-2003 epidemic and the SARS-CoV-2 from the current pandemic, from the MERS viruses of the 2012-2013 outbreak and from the strains related to the less lethal CoV OC43 (which uses MHC-class 1 molecules as a fusion receptor (26)) and bat strains related to HKU4 and HKU5. The MERS viruses, which use a different cellular receptor, the DPP4 protein, cluster with many camel isolates. The central nodes of the graph (red arrows) are viruses from bats that use the same DPP4 receptor as MERS (27).

**Figure 4.**
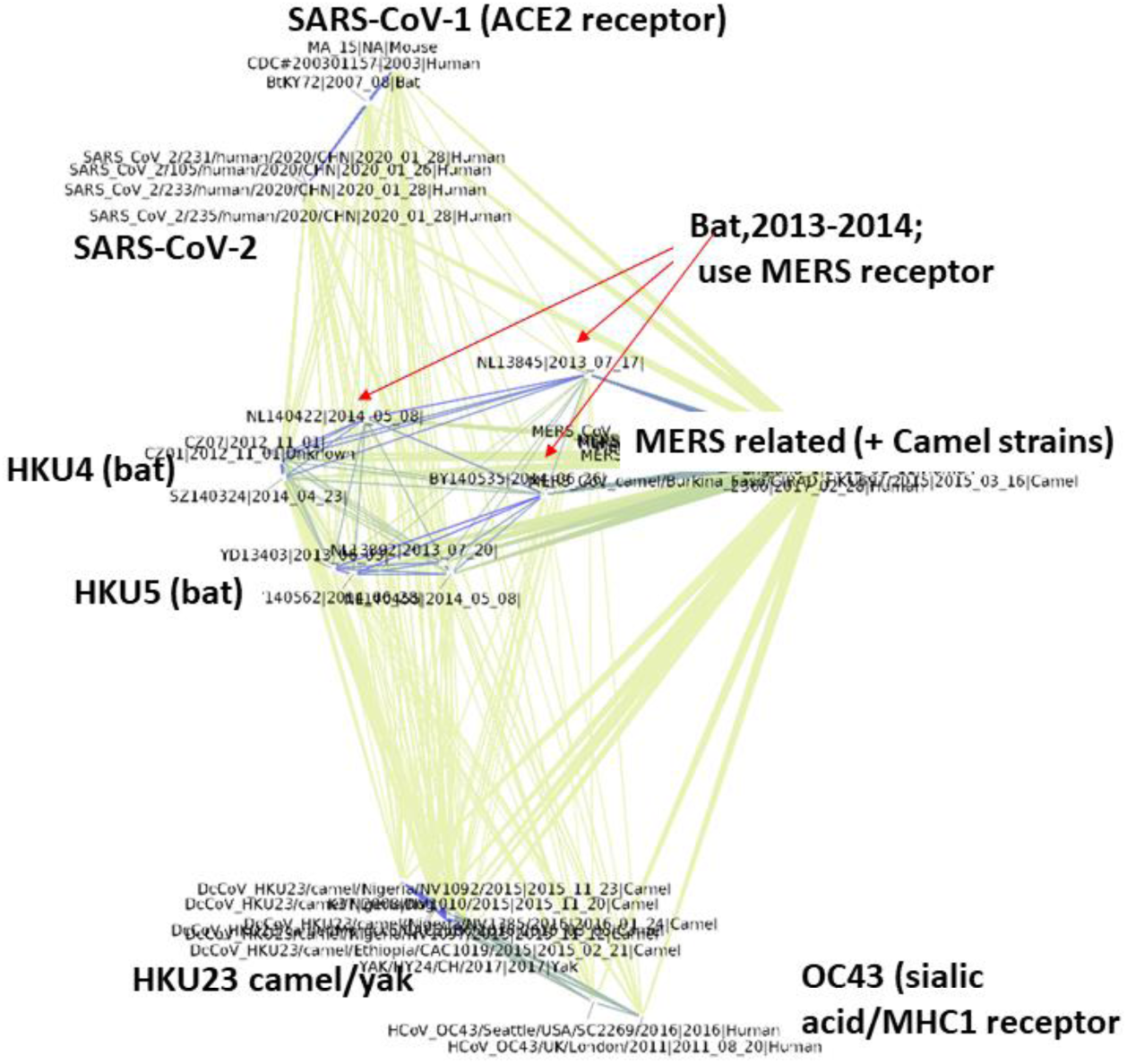
PD graph groups SARS-CoV-1 and -2 spike proteins and distinguishes them from other circulating β-CoV strains. The 2020 isolates of SARS-CoV-2, which like the SARS viruses use the human ACE2 receptor are closest to human SARS 2003, MA-15, from a human case in 2003 passaged in mice) and a 2007 strain from a bat than they are to other circulating β-CoV strains. Annotations indicate grouping according to receptor type, where known. Red arrows show nodes, bat viruses BY140535 (HKU5 related), NL13845 and NL140422 strains that use the MERS receptor, DPP4, all sequenced in China in 2013/2014. Blue lines show PD <7 (low PD= more similar), other lines are PD<14 whereby the thickness indicates the degree of relatedness.

The viruses closest to the SARS-CoV-2 sequences (from the current pandemic) are human SARS sequenced in 2003, M15, an isolate from a fatal human SARS infection that was passaged in mice and sequenced in 2003 at the NIH and a bat virus isolated in 2007 in China. The graph emphasizes the similarity of the two SARS viruses, which use the human ACE2 receptor for cell entry. It also suggests that SARS did not disappear in 2003, but rather continued to circulate in some mammalian vector for another 16 years, before reappearing in a much more contagious form to attack humans again.

## Discussion

Most phylogenetic analysis of viral sequences start with multiple sequence alignments. Aligning very diverse sequences, especially those containing multiple repeats or insertions, can be quite difficult (28). D-graph, as a flexible sequence analysis tool for protein sequences, provides a valuable first step to obtain an overview of related viral species without the need for an alignment. DGraph is unique in that it implements the PCP descriptors and PD calculations which were previously validated in our work in comparing allergenic proteins and their epitopes (6, 29, 30). No hypothesis about an ancestral sequence is required to follow inter-strain differentiation. As Fig. 1 illustrates, simply starting with a list of unaligned sequences in FASTA format, one can rapidly determine the interrelatedness of sequence data from two different virus families, the FV and EV (12), whereby the PD approach can give more detailed clustering than simple Clustal scores. As Figure 2-4 shows, automatic calculation of the PD values between even large groups of viral sequences yields PD-graphs consistent with what is known of the their vector, host range, receptor type (particularly for the β-CoV, Fig. 4) and disease phenotypes.

There are many methods being developed to handle and interpret the large amounts of sequence data available for viruses (31-33). A well done viral phylogeny is useful for suggesting the evolutionary relationships between viruses, their rates of change (34), and may also alert one to tipping points where additional changes may result in significant phenotypic variation (35) or viral outbreak (36, 37). Determining the interrelatedness of virus sequences is perhaps most important for the design of wide spectrum vaccines and treatments for viral diseases (7).

As shown here, D-graph can quantitatively present the interrelatedness of aligned sequences using established metrics such as Clustal scores. As such, it resembles other commonly used approaches for visualization of network connectivity, such as BioLayout (38), or Cytoscape (39, 40). However, the program’s default mode generates PD graphs even of large numbers of unaligned sequences (such as the >300 used for Figure 4). While other methods can graphically present protein sequences in an alignment free manner using numerical descriptors for the amino acids, and display them as connecting vectors in a curve in a 2D space (41, 42), they are most useful for comparing a few sequences. The D-graph program works from a list of unaligned sequences and can also use additional data, while allowing the user to adjust the program parameters to obtain results even for very distantly related sequences.

### PD-graphs suggest evolutionary paths between distantly related viruses

PD-graphs emphasize the non-linearity of viral evolution that may be missed when using linear phylogenetic trees to model the evolution of viral groups whereby there is no implied directionality in the connecting lines, which only represent the mathematical relationship between pairs. However, the PD-graph analysis of the FV (Fig. 2) has features that correlate with suggested paths for the divergence of the mammalian pathogens. The clustering emphasizes the central position of YFV, relative to both mosquito and tick born viruses and the “mosquito only” viruses.

This implies that the ability to circulate within their arthropod vectors may have taken priority during evolution of the mammalian pathogens. This may account for the relative stability of the YFV genome compared to that of the DENV types, which must have evolved under pressure from mammalian immune responses (34). Another interesting feature of the PD-graph is the grouping according to known host even when isolated from geographically very distant places. Entebbe (node 23) and Yokose (node 24) viruses, which were isolated from bats in Uganda (ENTV) (43) and Japan (YOKV) group together, but also have strong connectivity to other mosquito borne viruses. The YOKV sequence cross reacts with antibodies in sera from humans infected with DENV or after vaccination with YFV (35) and, depending on the protein area chosen, is similar to many different mosquito borne FV. Some of the early reports were not conclusive about the cross-reactivity of ENTV with other FV (44, 45). PD-graphs could provide testable hypotheses on FV cross-recognition, by comparing graphs made using inter-strain ELISA values and PD values.

YF in turn has connectivity to both the tick and mosquito borne viruses, which subdivide into well separated clusters according to their disease phenotype. The close relationship of the tick born viruses to one another suggests that their ancestry is relatively recent, or that other factors in the tick life cycle may constrain their evolution rate (46-48). The distinct properties of the tick vs the mosquito born viruses (49, 50) illustrated by their neatly defined clusters, reflects these influences.

## Conclusion

The D-graph program can be used to plot the interrelatedness of sequences according to physicochemical property similarity and suggest evolutionary relationships, without needing an alignment or assuming a common ancestor. While the figures shown here illustrate the application to viral proteins, any group of sequences or numerical relationships of objects, including immunological metrics, can in principle be used as input to the program. We thus anticipate that it will find numerous uses for the increasing numbers of virus sequences, as well as those for many other areas and are herewith releasing a downloadable version of the program.

## Acknowledgements

### Funding

This work was supported by grants from the National Institutes of Health [AI137332 to WB, AI064913 to WB/CHS, AI105985 to CHS]. The computational resources of the Sealy Center for Structural Biology and Molecular Biophysics were also used in this project.

