## Supplemental Material for "D-graph clusters flaviviruses and β-coronaviruses according to their hosts, disease type and human cell receptors"

**Supplementary Material**

**Table S1 List of the 49 flaviruses mapped by DGraph in fig.1 and fig.2.**

| ID | Flavi track ID | Genbank | Viral name | Length of Polyproteins |
|---|---|---|---|---|
| 1 | >CFAx92USxXmX | AIM49245 | Cell fusing agent virus (CFAV) | 3341 aa |
| 2 | >KRx99KExNmX_r | NP_891560 | Kamiti River virus | 3357 aa |
| 3 | >CX_NC_008604 | YP_899469 | Culex flavivirus | 3362 aa |
| 4 | >BSQx04USmXxX | NC_009026 | Aroa virus | 3429 aa |
| 5 | >IGUx04USmXxX | NC_009026.2 | Aroa virus (Iguape) | 3416 aa |
| 6 | >KOKx04USmXxX | AAV34157 | Kokobera virus | 3410 aa |
| 7 | >ALFx66AUmXxX | AAX82481 | Alfuy virus | 3434 aa |
| 8 | >MVEe99AUmXxX | NP_051124 | Murray Valley encephalitic virus | 3434 aa |
| 9 | >JBEe82JPmNmX_r | NP_059434 | Japanese encephalitis virus | 3432 aa |
| 10 | >USUx01ATmNiX_r | YP_164264 | Usutu virus | 3434 aa |
| 11 | >KUNx60AUmXmX | AAP78942 | Kunjin virus | 3433 aa |
| 12 | >WNe93DEmXxX_r | NP_041724 | West Nile virus (WNV) | 3430 aa |
| 13 | >SLEe04USmXxX_r | YP_001008348 | Saint Louis encephalitis virus | 3430 aa |
| 14 | >ILHx04USmXxX | YP_001040006 | Ilheus virus | 3424 aa |
| 15 | >ROCx04USmXxX | ATG32103 | Rocio virus | 3425 aa |
| 16 | >BAGx04USmXxX | AAV34161 | Bagaza virus | 3426 aa |
| 17 | >KEDx04USmXxX | AAV34156 | Kedougou virus | 3408 aa |
| 18 | >ZIKAx04USmXxX | YP_002790881 | Zika virus | 3419 aa |
| 19 | >DV1v74AUmNxX_r | NP_059433 | Dengue virus 1 | 3392 aa |
| 20 | >DV3x56PHmXxX_r | YP_001621843 | Dengue virus 3 | 3390 aa |
| 21 | >DV2v88USmXxX_r | NP_056776 | Dengue virus 2 | 3391 aa |
| 22 | >DV4v00USmXxX_r | NP_073286 | Dengue virus 4 | 3387 aa |
| 23 | >ENTx06UGxXrX | AKP24039 | Entebbe bat virus | 3411 aa |
| 24 | >YOKx71JPxNbX_r | NP_872627 | Yokose virus | 3425 aa |
| 25 | >SEPx06PGxXrX | YP_950478 | Sepik virus | 3405 aa |
| 26 | >YFx27GHmXhX | NP_041726 | Yellow fever virus | 3411 aa |
| 27 | >AHFh95SAtXhX_r | AFV78154 | Alkhumra hemorrhagic fever virus | 3416 aa |
| 28 | >KFDx03FRtXxX | YP_009513189 | Kyasanur Forest disease virus | 3416 aa |
| 29 | >GGE_DQ235153 | ABB90677 | Greek goat encephalitis virus | 3414 aa |
| 30 | >TSE_DQ235151 | ABB90675 | Turkish sheep encephalitic virus | 3414 aa |
| 31 | >LIx96GBtNtX_r | AYG86395 | Louping ill virus | 3414 aa |
| 33 | >SSE_DQ235152 | ABB90676 | Spanish sheep encephalitis virus | 3414 aa |
| 33 | >TBEe95ATtXrX | NP_043135 | Tick-borne encephalitis virus | 3414 aa |
| 34 | OMSKh02UStNtX | NP_878909 | Omsk hemorrhagic fever virus | 3414 aa |
| 35 | >LGTx56MYtXtX_r | NP_620108 | Langat virus | 3414 aa |
| 36 | >DTx95CTAtXxX | AAL32169 | Deer tick virus | 3415 aa |

| | | | | |
|---|---|---|---|---|
| 37 | >POWx58CAtXxX_r | NP_620099 | Powassan virus | 3415 aa |
| 38 | >GGY_DQ235145 | YP_009345034 | Gadgets Gully virus | 3416 aa |
| 39 | >KSIx04UZtXxX_r | ABE73208 | Karshi virus | 3416 aa |
| 40 | >RF_DQ235149 | ABB90673 | Royal Farm virus | 3417 aa |
| 41 | >KAD_DQ235146 | ABB90670 | Kadam virus | 3404 aa |
| 42 | >MEA_DQ235144 | ABB90668 | Meaban virus | 3421 aa |
| 43 | >SRE_DQ235150 | ABB90674 | Saumarez Reef virus | 3422 aa |
| 44 | >TYU_DQ235148 | ABB90672 | Tyuleniy virus | 3422 aa |
| 45 | >MMLe58USxNbX | NP_689391 | Montana myotis leukoenc. virus | 3374 aa |
| 46 | >RBx99FRxXxX_r | NP_620044 | Rio Bravo virus | 3379 aa |
| 47 | >MODx58USxNrX | NP_619758 | Modoc virus | 3374 aa |
| 48 | >APOIX99FRxNrX | NP_620045 | Apoi virus | 3371 aa |
| 49 | >TABx73TTxXbX | NP_658908 | Tamana bat virus | 3350 aa |

**Fig. S1: PD graph of the E domain 3 of sequences representing WNV, the four DENV serotypes, and Zika virus.** The 7P8 sequence is a PCP-consensus derived from DENV strains that represents all four serotypes of DENV. Note that Zika has only slightly higher identity to WNV than it does to the DENV consensus.

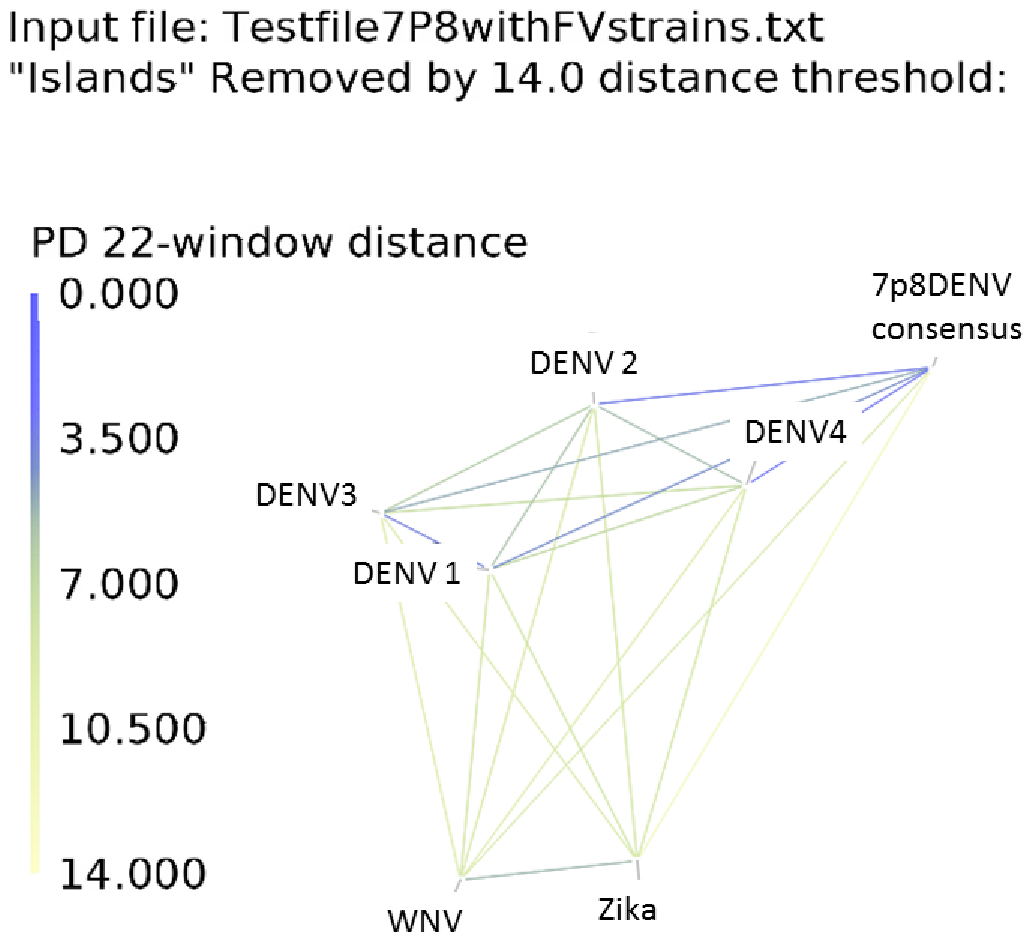

**Using the program**: First time users should follow directions in the “read me” file. Input files should be ASCI text files. The headers of each FASTA sequence file can directly be displayed on the resulting graph by pushing “y” to stop the program and “b” to show sequence headers. The input files for the figures in this paper can be obtained from the GitHub website.

To open the program, click on the executable .jar file. The program will automatically open and supply a list of subroutines for formatting or inputting data. In default mode, where the user supplies a list of sequences in FASTA format, the program will automatically open the selected text file, calculate the inter-sequence PD values as described below, and begin the simulation to determine the best 2D-presentation of the data. For very diverse data, it is recommended that the user scale the viewing field by pushing 9 to zoom out (this may require pushing 9 several times to get the data into view). After the viewing field is established, the points can be rescattered by pushing ‘e”, at which point the simulation will start again.  The user can stop the simulation or restart it at any time by pushing “y’. Once the user is satisfied with the graph, it can be saved as a .pdf and .png file by typing s.
